# Vagus nerve stimulation by focused ultrasound attenuates acute myocardial ischemia/reperfusion injury predominantly through cholinergic anti-inflammatory pathway

**DOI:** 10.1101/2023.12.05.570312

**Authors:** Qian Zhang, Qianyun Cai, Shenrong Zhong, Qin Li, Weibao Qiu, Juefei Wu

**Author notes:** Correspondence to Juefei Wu, Department of Cardiology, Nanfang Hospital, Southern Medical University, No. 1838, North of Guangzhou Avenue, Guangzhou, China. 510515.

## Abstract

**Background:** The invasive method using electrically stimulating vagus nerve has been demonstrated to inhibit the inflammatory response against acute myocardial ischemia/reperfusion(I/R) injury. This neuro-regulatory technique used is invasive implantation of electrical stimulators which have the risk of infection and other complications. A non-invasive method of vagus nerve stimulation applying on acute myocardial I/R injury is particularly important and desirable.

**Objectives:** This study aimed to test the hypothesis that focused ultrasound stimulation(FUS) of vagus nerve attenuates acute myocardial I/R injury through cholinergic anti-inflammatory pathway.

**Methods:** This study was completed with two parts. In the first part, ischemia was induced by left anterior descending coronary artery (LAD) occlusion for 30 minutes (min), followed by 2 hours (h) of reperfusion, FUS lasted for 50 min after reperfusion for 10 min. Infammatory markers, arrhythmia score, histopathology, infammatory markers α7 nicotinic acetylcholine receptor (α7nAchR) protein were determined. In the second part, ischemia was induced by LAD occlusion for 30 min, followed by 24h of reperfusion, FUS lasted for 50 min after reperfusion for 10 min. Myocardial fibrosis, infarct size, echocardiographic measurements, biomarkers of pro-oxidative stress, antioxidant and myocardial apoptosis were determined.

**Results:** FUS of vagus nerve significantly decreased the level of infammatory markers in I/R+FUS group compared with the I/R group, I/R+vagotomy group and I/R+vagotomy+FUS group. FUS decreased the incidence of ventricular tachycardia (VT) and ventricular fibrillation (VF), while also reduced arrhythmia score in I/R+FUS group compared with the I/R group. These improvements were associated with the inhibition of acute inflammatory reaction through cholinergic anti-inflammatory pathway. FUS limited myocardial fibrosis and infarct size, preserved left ventricular function. This cardioprotective effect was associated with alleviation of apoptosis and oxidative stress.

**Conclusions:** The noninvasive method of ultrasonic neuromodulation using FUS of vagus nerve exerted protective effects on acute myocardial I/R injury. Its potential mechanisms involved the suppression of acute inflammatory reaction through cholinergic anti-inflammatory pathway.

## Introduction

Coronary heart disease is a common cardiovascular disease, which is a severe hazard to people’s health. Acute myocardial infarction is one of the main causes of death of coronary heart disease. The rapid and effective reperfusion to restore coronary blood flow is the most effective way to save viable cardiomyocytes and reduce the morbidity and mortality of patients with myocardial infarction. However, previous results^[1,2]^ have shown that the damaged myocardial function does not recover after the flow of coronary artery is re-established, in fact, the myocardial injury is aggravated, and even the infarct area is expanded. This is called myocardial I/R injury, which is also considered to be a “double-edged sword”.

The mechanism of myocardial I/R injury is very complex. Acute inflammatory reaction is the initiating factor of myocardial I/R injury. After myocardial I/R, a variety of cells in the blood gather, adhere and activate, and release a large amount of inflammatory factors, such as tumor necrosis factor (TNF-α), interleukin-1 (IL-1) and interleukin-6 (IL-6) causing cardiomyocyte injury. Cardiomyocyte injury leads to myocardial energy metabolism disorder and microcirculation disorder, which further aggravates apoptosis and inflammatory reaction. Because inflammatory reaction plays a very important pathogenic role in the occurrence and development of myocardial I/R injury, myocardial I/R injury can thus be alleviated by regulating inflammatory reactions. Previous studies have shown that stimulation of peripheral vagus nerve and application of cholinergic transmitter acetylcholinergic (Ach) can inhibit systemic inflammatory response syndrome and significantly reduce the release of cytokines^[3]^. This pathway is cholinergic anti-inflammatory pathway. Studies have confirmed that activating cholinergic anti-inflammatory pathway by electrically stimulating vagus nerve can reduce the level of proinflammatory cytokines in circulating blood and inhibits systemic inflammatory response^[4]^. Previous studies have also shown that focused ultrasound is a good non-invasive way of stimulation in vitro^[5]^. We hypothesized that ultrasound stimulation of vagus nerve can activate cholinergic pathway, inhibits inflammatory response, protects cardiomyocytes and reduces myocardial I/R injury. This study will construct a rat model of acute myocardial I/R injury, observes the effect of vagus nerve stimulated by focused ultrasound on acute myocardial I/R injury and its mechanism.

## Methods

### Animal preparation

All experiments were approved by the Institutional Animal Care and Use Committees of the Nanfang Hospital, Southern Medical University, China. Seventy two 8 weeks male Sprague-Dawley(SD) rats (160–230g) were anesthetized by an intraperitoneal injection of 50mg/kg pentobarbital sodium. Trachea cannula was inserted and was connected to the respirator. Surface electrocardiogram (lead II) and heart rate (HR) were monitored, and all data were recorded for subsequent analysis.

### Experimental groups

In the first part, forty five SD rats were randomly divided into five groups: Sham group (N=9): the LAD was not ligatured after thoracotomy. I/R group (N=9): the LAD was ligatured for 30 min, followed by reperfusion for 2h. I/R+vagotomy group (N=9): bilateral cervical vagus nerve vagotomy, the LAD was ligatured for 30 min, followed by reperfusion for 2h. I/R+vagotomy+FUS group (N=9): bilateral cervical vagus nerve vagotomy, the LAD was ligatured for 30 min, followed by reperfusion for 2h, the right cervical vagus nerves were stimulated by focused ultrasound for 50 min after reperfusion for 10 min. I/R+FUS group (N=9): the LAD was ligatured for 30 min, followed by reperfusion for 2h, the right cervical vagus nerves were stimulated by focused ultrasound for 50 min after reperfusion for 10 min. Each group was divided into three subgroups (N=3). The hearts of experimental rats in the three subgroup were respectively used for inflammatory factors detection, H&E staining and detection of expression of α7nAchR.

In the second part, twenty seven SD rats were randomly divided into three groups: Sham-24h group (N=9, the LAD was not ligatured after thoracotomy), I/R-24h group (N=9, LAD was occluded for 30 min followed by reperfusion for 24h), and I/R-24h+FUS group (N=9, LAD was occluded for 30 min followed by reperfusion for 24h, FUS lasted for 50 min after reperfusion for 10 min). Each group was divided into three subgroups (N=3). The hearts of experimental rats in the three subgroup were respectively used for Masson staning, detection of biomarkers of pro-oxidative stress, antioxidant and myocardial apoptosis, and staining with Evans blue and triphenyltetrazolium chloride (TTC).

### Acute myocardial I/R injury protocol

Left thoracotomy was performed and the LAD coronary artery exposed. The LAD was ligatured to produce an acute I/R injury model. Occlusion was achieved via 5-0 silk suture tie and lasted for 30 min. Then the LAD ligature was released and the arterial circulation was re-established for 2h or 24h. The standard of successful ischemia included that: the local cyanosis occurred in the myocardial ischemic area, the ECG of lead II showed the ST segment elevation significantly. The standard of successful reperfusion included: the part of the ischemic area turned red and lead II of ECG showed leveling of the ST segments.

### Focused ultrasound stimulation on vagus nerve

Focused ultrasound stimulator was located on the right neck of the rats. The following stimulation parameters (Fig. 1A) were explored by a stimulator (Ultrasound Neurostimulation System, Shenzhen Institutes of Advanced Technology, Chinese Academy of Sciences, China), Frequency: 1.71MHz, Amplitude: 50per, T1: 10ms, T2: 100ms, T3: 360s, Toff: 4 minutes. The stimulation lasted for 50 min after reperfusion for 10 min.

**Figure 1.**
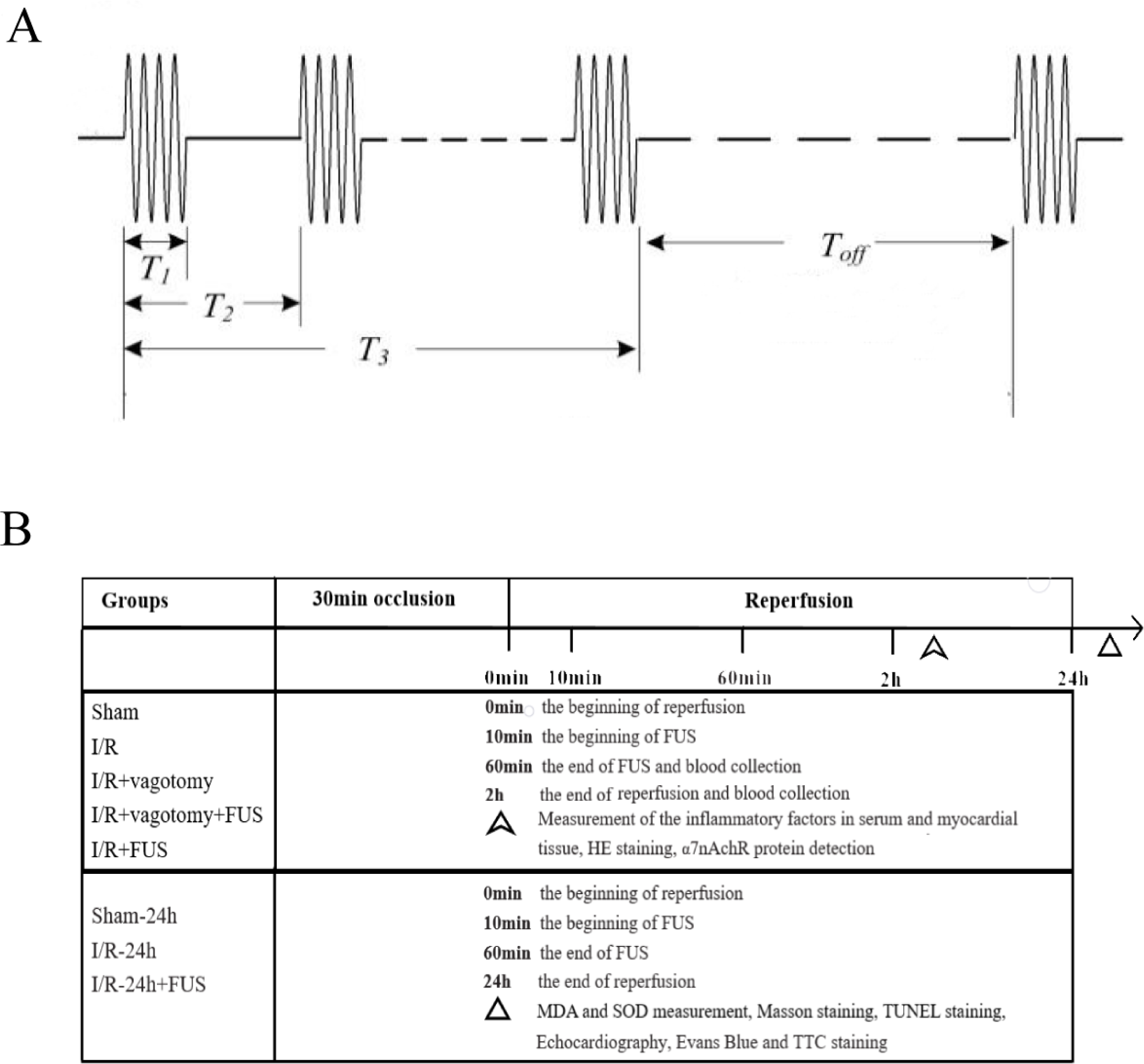
Experimental protocol. (A) Focused ultrasound stimulation parameters. T1: continuous pulse duration, T2: pulse repetition period, T3: stimulation duration, Toff: off time. (B) Experimental protocol showing the groups, intervention and outcomes measurement.

### Measurement of ventricular arrhythmias incidences

In the first part, electrocardiogram (ECG) was continuously recorded to detect the incidence and duration of ventricular arrhythmias. According to the Lambeth diagnostic convention criteria^[6]^ with modifications, ventricular arrhythmias were classified as premature ventricular contraction (PVC), VT and VF. Specifically, PVC was defined as ventricular contractions without atrial depolarization, whereas VT was characterized by more than six consecutive PVCs. VF was defined as a loss of synchronicity of the ECG. Furthermore, the arrhythmia scores all correlated with the incidences of PVC, VT and VF. Arrhythmia scores were tabulated for the entire 30 min ischemia period and 120 min reperfusion period using Score A as described previously by Curtis and Walker^[7]^. Briefly, each heart was given a score based on the following criteria: 0: <50 ventricular premature beats, 1: 50-499 ventricular premature beats, 2: >500 ventricular premature beats and/or one episode of spontaneously reverting ventricular tachycardia or ventricular fibrillation, 3: more than one episode of spontaneously reverting ventricular tachycardia or fibrillation (<1 min total combined duration), 4: 1-2 min of total combined ventricular tachycardia or fibrillation, 5: >2 min of ventricular tachycardia or fibrillation.

### Measurement of the inflammatory factors in serum and ischemic myocardial tissue

In the first part, blood was collected through the right jugular veins of the experimental animals after thoracotomy at 90 min and 150 min in sham group, and after reperfusion at 60 min and 120 min in other four groups, respectively. Blood samples sat under room temperature for 30 min and then were centrifuged at 3,000 r/min for 10 min. The supernatant was taken and placed in the EP tube. The ischemic myocardial tissue were centrifuged and the supernatant were taken and placed in the EP tube. All serum and myocardial specimen were stored in the refrigerator under −80°C. The enzyme-linked immunosorbent assay (ELISA) kits (Meimian Industrial Co., Ltd, Jiangsu, China) for rats was used to detect the levels of C-reactive protein (CRP), IL-1, IL-6, high mobility group protein 1 (HMGB-1), cardiac troponin T (Tn-T), intercellular adhesion molecule-1 (ICAM-1), Ach, TNF-α in the serum, and the levels of IL-1, Tn-T, TNF-α in the ischemic myocardial tissue.

### Observation of neutrophil infiltration with Hematoxylin and Eosin (HE) staining

In the first part, the myocardial tissue were preserved with 4% paraform. Through the conventional dehydration, wax infiltration and embedding, The myocardial tissue below ligature line were made into paraffin section to be stained with HE. And then the situation of neutrophil granulocyte infiltration was observed under optical microscope (×100).

### α7nAchR protein expression detection

In the first part, α7nAchR protein of the myocardial tissue was stained with brown positive granules. According to the degree of color, they were divided into three types, no brown granules was negative, light brown granules was positive, dark brown granules was strong positive. Image Pro Plus 6.0 software was used to select brown positive granules and analyzed immunohistochemical cumulative optical density value (IOD) of each group.

### Echocardiographic measurements

In the second part, myocardial function was accessed by echocardiography at 24h after reperfusion using the ultrasound system with the MS250 probe (Vevo 2100, Visual sonics, USA). Short-axis B-mode images of the left ventricle were acquired at the level of the papillary muscles. Left ventricular end-systolic diameter (ESd), end-diastolic diameter (EDd), left ventricular fractional shortening (LVFS) and left ventricular ejection fraction (LVEF) were measured.

### Observation of myocardial fibrosis with Masson staining

Masson staining were performed to analyze myocardial fibrosis. Myocardial tissues were preserved with 4% paraform. Thereafter, paraffin-embedded sections were stained with Masson, and observed under microscope.

### Ischemic and infarct size determination

In the second part, the myocardial infarct size was assessed with 0.5% Evans Blue and 1.0% TTC staining. After 24h of reperfusion, the descending aorta was clamped with a vascular clamp. Evans blue solution was infused into ascending aorta, which resulted in a dark blue staining of the non-ischemic area. Then, the heart was rapidly excised, removed, and irrigated with normal saline to wash out blood from chambers and vessels. Removed heart was cut into 4 mm coronal slices. Slices were placed in the vital dye TTC at 37 °C in the dark for 30 min. Non-ischemic myocardium was stained blue with Evans blue and ischemic myocardium was stained red with TTC, whereas infarct myocardium was not stained blue or red but appeared white. The infarct size (white) and the area at risk (AAR) (red) from the section on the mitral side were determined by using image tool software Image-J. The area ratio of infarct size to the AAR and of the AAR to the ventricular area was calculated and expressed as a percentage.

### Determination of myocardial apoptosis

In the second experimental period, myocardial apoptosis was determined by TUNEL staining. The apoptotic cells were stained brown. The number of TUNEL positive cells and the total number of nuclei cardiomyocyte per high-powered field were counted using Image-J (National Institutes of Health, USA) from at least 6 randomly selected fields from the area at risk (AAR) in each section. All measurements were performed in a blinded manner.

### Measurement of MDA and SOD content in myocardium

In the second part, after 24h of reperfusion, myocardial tissue was obtained and homogenated with appropriate buffer, the supernatant was collected and stored at −80℃ after centrifugation. Superoxide dismutase (SOD) and malondialdehyde (MDA) were measured using colorimetry (Shenzhen TopBiotech Co., Ltd. Guangdong, China).

### Statistical analysis

Statistical analyses were performed using version 25.0 software (SPSS, Inc., Chicago, IL, USA). The data were tested for normality and homogeneity of variance. If it conformed to the normal distribution and homogeneity of variance, analysis of variance was used. Bonferroni was used for pairwise comparison between groups. Otherwise it was expressed as median (interquartile spacing), and Kruskal-Wallis H rank sum test was used. Data distribution of each group was drawn using GraphPad Prism 8.0.2. P<0.05 was considered to indicate a statistically significant difference.

## Results

### ECG parameters during I/R

The LAD occlusion could cause local cyanosis in the myocardial ischemic area and the synchronous lead II on ECG showed the ST segment elevation significantly. 30 min of LAD occlusion, followed by 120 min of reperfusion resulted in some part of the ischemic area became red and the synchronous lead II on ECG showed the ST segment leveling on reperfusion period. Fig. 2A shows examples of ECG tracing at baseline, ischemia and reperfusion. Ventricular arrhythmia occurred in both ischemia and reperfusion periods. Representative tracings of PVC, VT, and VF are shown in Fig. 2A.

**Figure 2.**
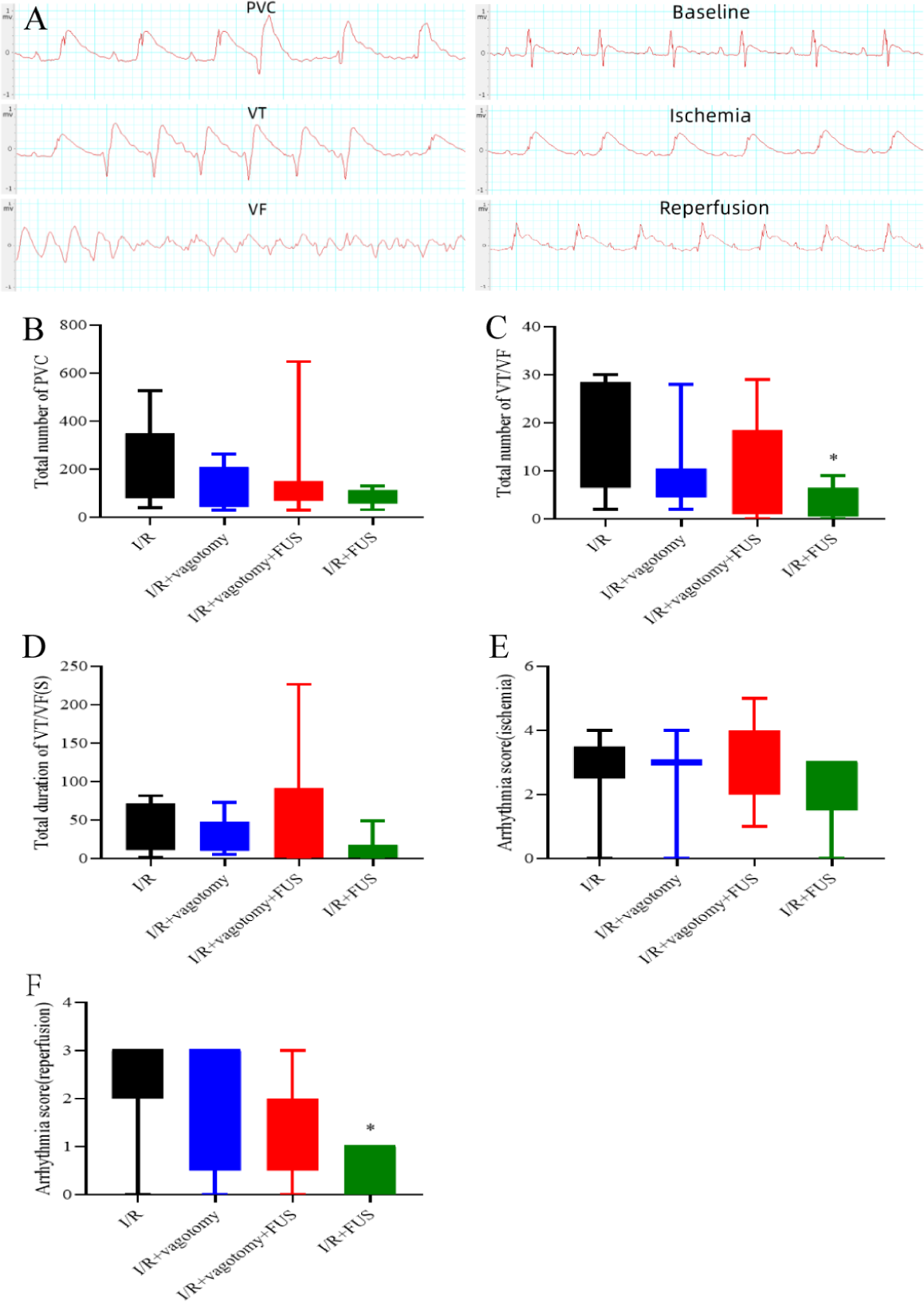
FUS decreased the incidence of ventricular arrhythmia. (A) Representative ECG tracings at the baseline, ischemia and reperfusion. Representative morphology of the PVC, VT and VF. (B) The total number of PVC. (C) The total number of VT/VF. (D) The duration of VT/VF. (E) The arrhythmia score during ischemia period. (F) The arrhythmia score during reperfusion period. * *P* < 0.05 vs. I/R group. ^†^ *P* < 0.05 vs. I/R+vagotomy group. ^‡^ *P* < 0.05 vs. I/R+vagotomy+FUS group.

### FUS decreased the occurrence of ventricular arrhythmia during I/R

The total number of VT/VF episodes during I/R period and the arrhythmia score during reperfusion period were significantly reduced in I/R+FUS group compared with the I/R group (*P* < 0.05, Fig 2C and F). However, the total number of PVC, the total duration of VT/VF onset and the arrhythmia score during ischemia period were not significantly different among I/R group, I/R+vagotomy group, I/R+vagotomy+FUS group and I/R+FUS group (*P* > 0.05, Fig 2B, D and E). The total number of VT/VF episodes during I/R period and the arrhythmia score during reperfusion period in I/R+FUS group were not significantly different compared with the I/R+vagotomy group and I/R+vagotomy+FUS group (*P* > 0.05, Fig 2C and F).

### FUS alleviated the serum inflammatory factors

Compared with the sham group, the differences of IL-6, CRP, Tn-T, HMGB-1, ICAM-1, TNF-α and Ach had no statistical significance in the I/R+FUS group after reperfusion for 60 min and 120 min (*P* > 0.05, Fig 3B, C, D, E, F, G and H), while IL-1 increased after reperfusion for 60 min (*P* < 0.05, Fig 3A). However, IL-1, IL-6, CRP, Tn-T, HMGB-1, ICAM-1 increased while Ach decreased in the I/R, I/R+vagotomy and I/R+vagotomy+FUS group after reperfusion for 60 min and 120 min, and TNF-α increased after reperfusion for 120 min (*P* < 0.05, Fig 3A, B, C, D, E, F, G and H). IL-1, IL-6, CRP, Tn-T, HMGB-1, ICAM-1 and TNF-α in the I/R+FUS group markedly decreased after reperfusion for 60 min and 120 min compared with those in the I/R group, while the Ach increased (*P* < 0.05, Fig 3A, B, C, D, E, F, G and H). Compared with those in the I/R+FUS group, IL-1, CRP, Tn-T, HMGB-1, ICAM-1 and TNF-α increased while Ach decreased in the I/R+vagotomy and I/R+vagotomy+FUS groups after reperfusion for 60 min and 120 min (*P* < 0.05, Fig 3A, C, D, E, F, G and H), and IL-6 increased after reperfusion for 120 min (*P* < 0.05, Fig 3B).

**Figure 3.**
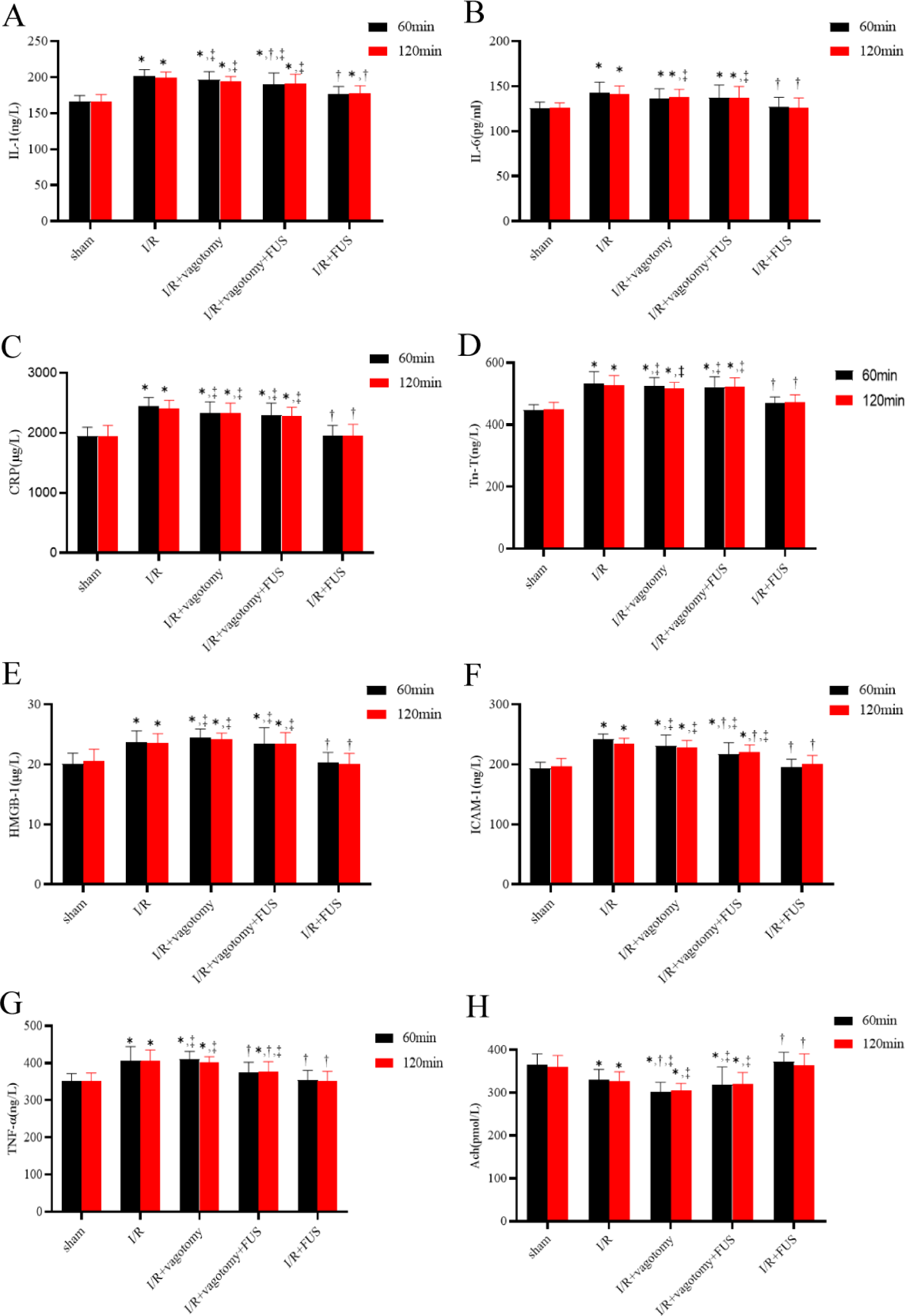
Effect of FUS treatment on inflammatory factors in the serum. (A) IL-1 levels in the serum. (B) IL-6 levels in the serum. (C) CRP levels in the serum. (D) Tn-T levels in the serum. (E) HMGB-1 levels in the serum. (F) ICAM-1 levels in the serum. (G) TNF-α levels in the serum. (H) Ach levels in the serum. * *P* < 0.05 vs. sham group. ^†^ *P* < 0.05 vs. I/R group. ^‡^ *P* < 0.05 vs. I/R+FUS group.

### FUS alleviated the myocardial inflammatory factors

Compared with the sham group, the levels of Tn-T, IL-1 and TNF-α in myocardium increased in the I/R, I/R+vagotomy and I/R+vagotomy+FUS groups (*P* < 0.05, Fig 4A, B and C), the differences of Tn-T, IL-1 and TNF-α had no statistical significance in the I/R+FUS group (*P* > 0.05, Fig 4A, B and C). Tn-T, IL-1 and TNF-α in the I/R+FUS group decreased compared with those in the I/R, I/R+vagotomy and I/R+vagotomy+FUS groups (*P* < 0.05, Fig 4A, B and C). Tn-T and TNF-α in the I/R+vagotomy group increased compared with those in the I/R group (*P* < 0.05, Fig 4A and C). IL-1 in the I/R+vagotomy+FUS group decreased compared with those in the I/R group (*P* < 0.05, Fig 4B).

**Figure 4.**
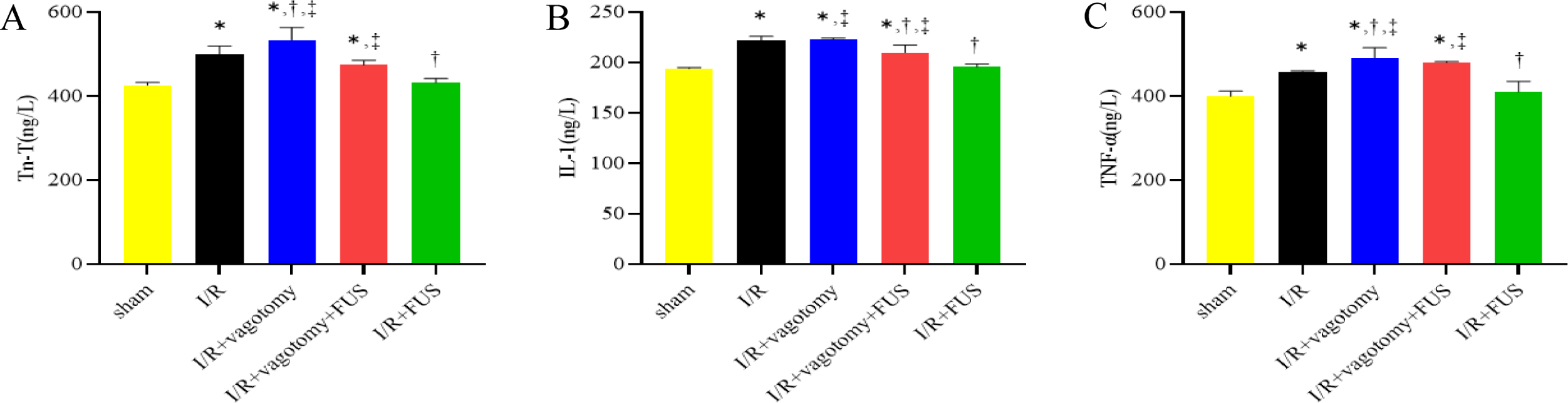
Effect of FUS treatment on inflammatory factors in the myocardium. (A) Tn-T levels in the myocardium. (B) IL-1 levels in the myocardium. (C) TNF-α levels in the myocardium. * *P* < 0.05 vs. sham group. ^†^ *P* < 0.05 vs. I/R group. ^‡^ *P* < 0.05 vs. I/R+FUS group.

### Observation of neutrophil infiltration with HE staining

A lot of neutrophil infiltration was seen in the I/R, I/R+vagotomy and I/R+vagotomy+FUS groups. In contrast, there was no obvious neutrophil infiltration noted in the sham and I/R+FUS groups(Fig. 5A).

**Figure 5.**
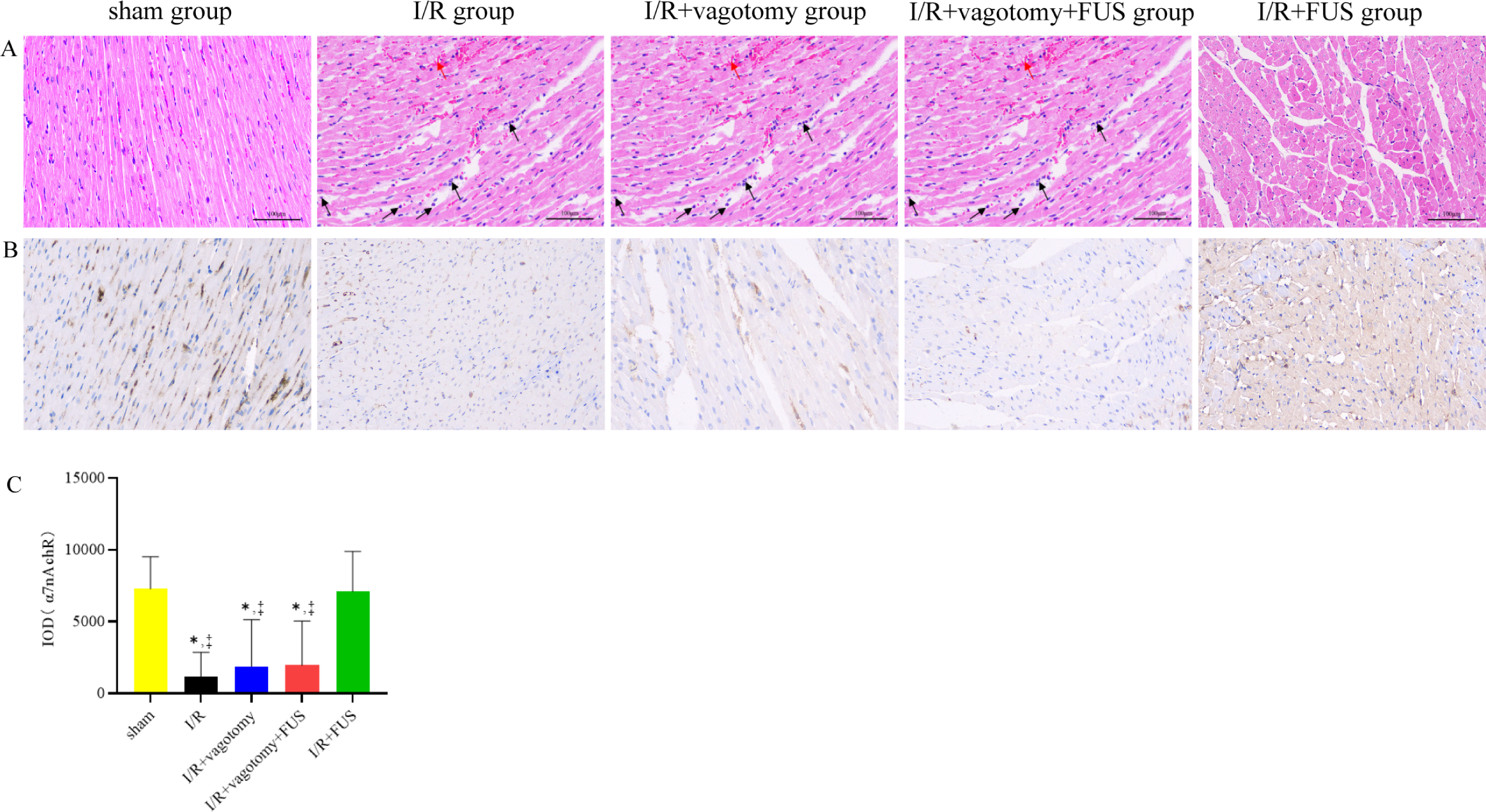
FUS attenuates I/R injury through cholinergic anti-inflammatory pathway. (A) Neutrophil infiltration of myocardial tissue in the sham, I/R, I/R+vagotomy, I/R+vagotomy+FUS and I/R+FUS groups (HE staining, ×200). The black arrows marked the neutrophil granulocyte. Cardiomyocytes were disordered and atrophic as indicated by red arrows. Yellow arrows marked interstitial bleeding. (B) Brown positive granules represent the positive protein expressions of α7nAchR in the myocardial tissue. (C) Comparison of α7nAchR protein expressions. \**P* < 0.05 vs sham group, ^‡^ *P* < 0.05 vs I/R+FUS.

### α7nAchR protein expression detection

The results of protein expression of the myocardial tissue showed that, brown positive granules were seen in the sham group and I/R+FUS group, while small amount of brownish positive granules was seen in the I/R, I/R+vagotomy and I/R+vagotomy+FUS groups(Fig. 5B). Comparing immunohistochemical cumulative optical density value (IOD) with those in the sham group, the α7nAchR protein expression in the I/R, I/R+vagotomy and I/R+vagotomy+FUS groups decreased obviously(*P* < 0.05), there was no statistical significance in I/R+FUS group. Compared with those in the I/R group, the α7nAchR protein expression in the I/R, I/R+vagotomy and I/R+vagotomy+FUS groups showed obvious decreased (*P* < 0.05, Fig. 5C).

### FUS preserved left ventricular function

Left ventricular function was evaluated by echocardiography 24 hours after reperfusion. The representative images show the short-axis view of the left ventricle in M-mode. Significant left ventricular dysfunction was observed in rats subjected to I/R-24h group compared with the sham-24h group. I/R-24h caused an increase in ESd, and decrease in LVEF and LVFS. LVEF and LVFS were improved in I/R-24h+FUS group compared with I/R-24h group, which shows the preservation of left ventricular systolic function(Fig. 6).

**Figure 6.**
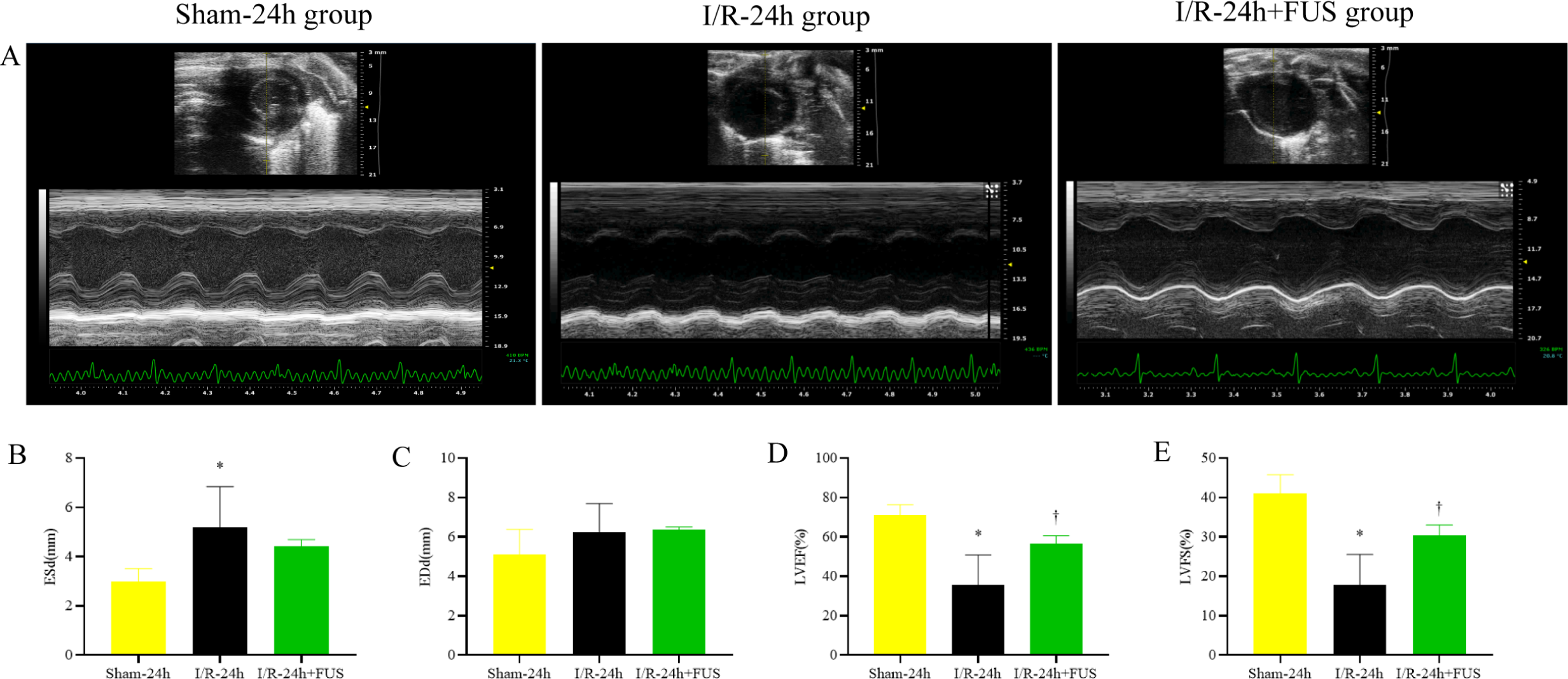
FUS preserved left ventricular function. (A) The representative echocardiography images of short-axis view of the left ventricle in M-mode. (B) Comparison of left ventricular ESd. (C) Comparison of left ventricular EDd. (D) Comparison of LVEF. (E) Comparison of LVFS. \**P* < 0.05 vs Sham-24h group, ^†^*P* < 0.05 vs I/R-24h group.

### FUS alleviated I/R induce myocardial fibrosis

Masson staining used to determine if the observed protective effect of FUS against I/R injury was associated with decreased fibrosis. The proportions of Masson staining were significantly increased in the I/R-24h group compared to the Sham-24h group, significantly decreased in the I/R-24h+FUS group compared with the I/R-24h group (Fig. 7A and B).

**Figure 7.**
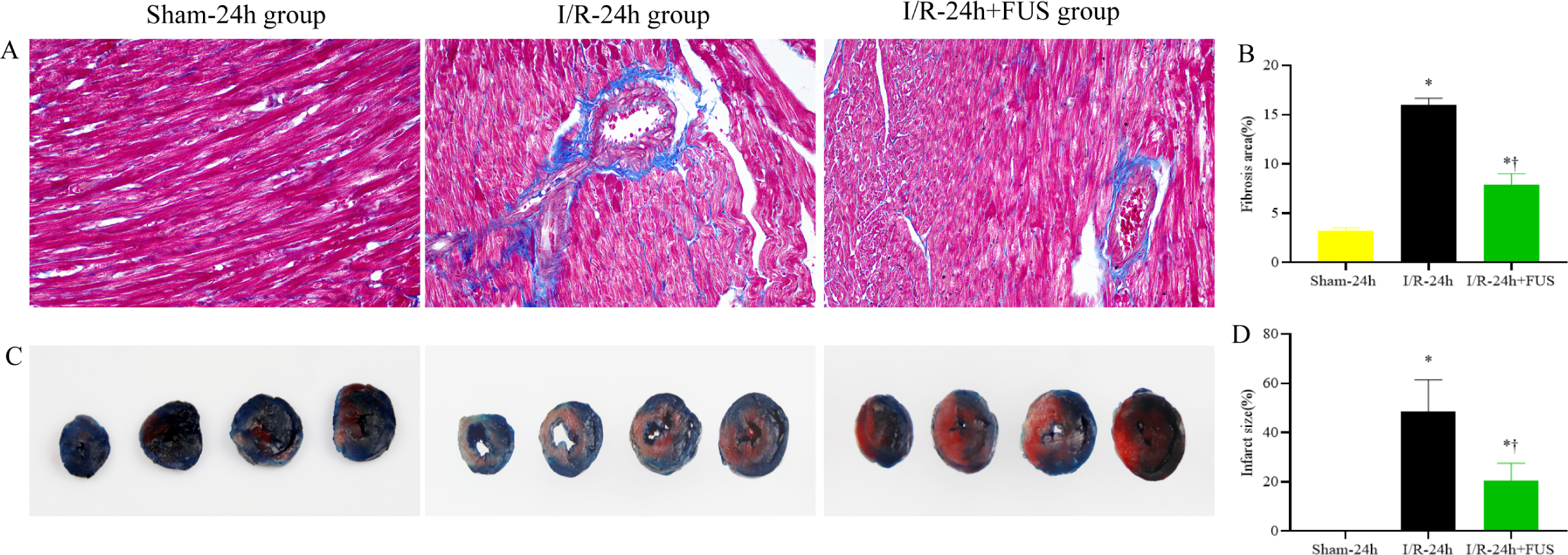
FUS limited the myocardial fibrosis and infarct size. (A) Representative pictures of sections stained with Masson (×200). Collagen fibers stained in blue indicated. (B) Comparison of the fibrosis area. (C) Representative photographs of mid-ventricular cross sections stained with Evans/TTC. Dark blue stain indicated viable area; White stain indicated infarct region; Red stain indicated area at risk. (D) Comparison of the infarct size. \**P* < 0.05 vs Sham-24h group, ^†^*P* < 0.05 vs I/R-24h group.

### FUS limited the myocardial ischemic and infarct size

As shown in Fig 5, myocardial infarct size was expressed as the percentage of AAR. The AAR was significantly reduced in the I/R-24h+FUS group than the I/R-24h group(Fig. 7C and D).

### FUS alleviated I/R induce apoptosis

Apoptosis plays a critical role in I/R. TUNEL staining was used to determine if the observed protective effect of FUS against I/R injury was associated with decreased apoptosis. In response to I/R, total TUNEL positive cells were significantly increased compared to the Sham-24h group. Significant reduction in TUNEL positive cells was noted in the I/R-24h+FUS group compared with the I/R-24h group(Fig. 8A and B).

**Figure 8.**
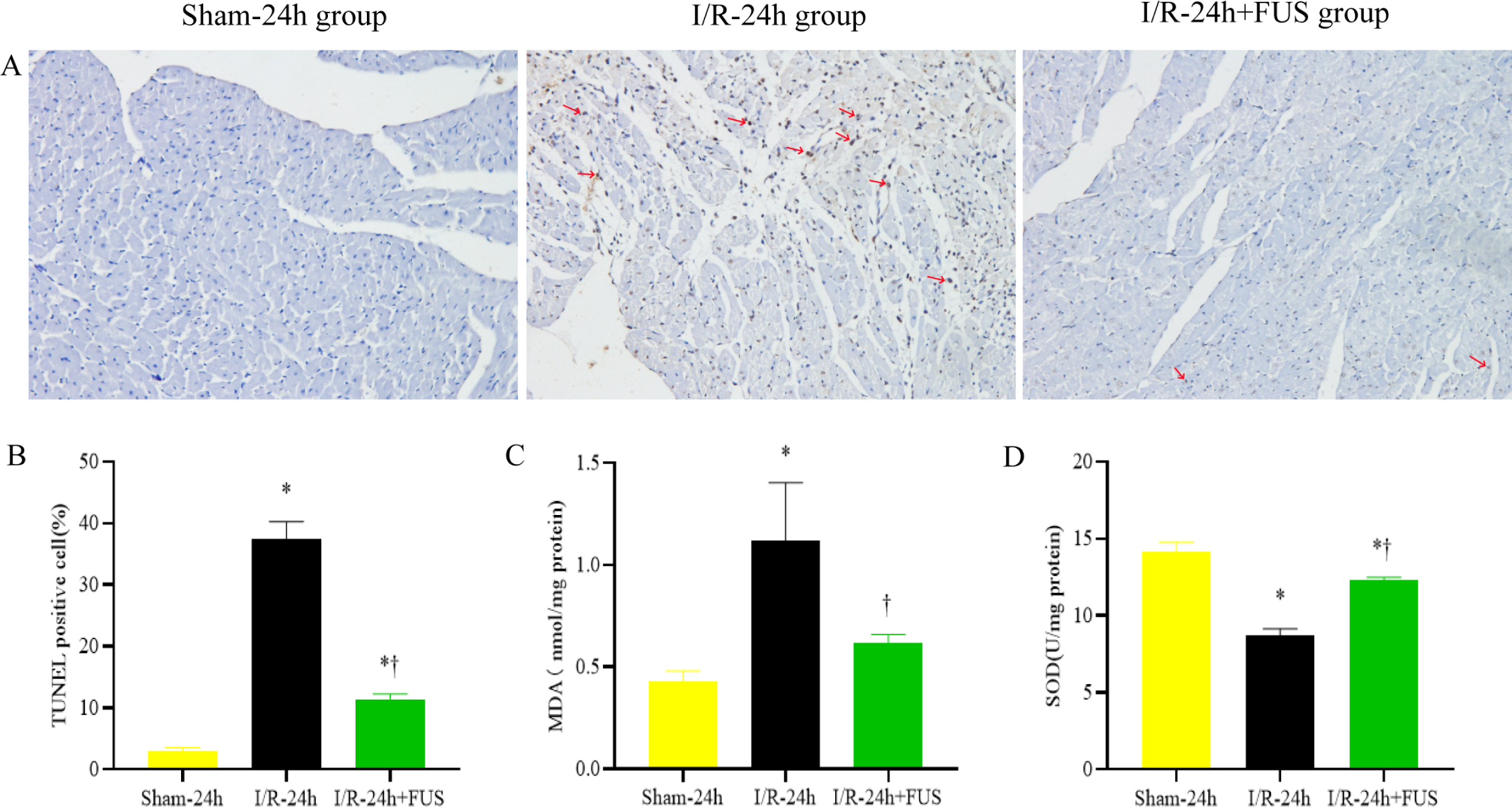
FUS alleviated *apoptosis and* oxidative stress. (A) Representative pictures of sections stained with TUNEL (×100). The red arrows marked brown staining indicated TUNEL positive apoptotic nuclei. (B) Comparison of the TUNEL positive cell. (C) MDA level in the myocardium. (D) SOD level in the myocardium. \**P* < 0.05 vs Sham-24h group, ^†^*P* < 0.05 vs I/R-24h group.

### FUS attenuated I/R induce oxidative stress

MDA and SOD were determined as a biomarker of pro-oxidative stress and antioxidant respectively. I/R significantly increased the MDA level and reduced the SOD level in myocardium in I/R-24h+FUS group compared with the Sham-24h group. There was a marked decreased in MDA and increased in SOD in I/R-24h+FUS group in comparison with the I/R-24h group(Fig. 8C and D).

## Discussion

### Main findings

Our results demonstrated that vagus nerve stimulation by focused ultrasound attenuates acute myocardial I/R injury by inhibiting inflammatory reaction through cholinergic anti-inflammatory pathway. FUS of vagus nerve significantly decreased the maker of myocardial necrosis Tn-T, and the inflammatory markers CRP, IL-1, IL-6, HMGB, ICAM-1 and TNF-α in the serum and decreased the levels of Tn-T, IL-1 and TNF-α in the myocardium. We also observed that FUS on vagus nerve in I/R+FUS group prevented the incidence of spontaneous VT/VF episodes during I/R period and reduced the arrhythmia score during reperfusion period compared with the I/R group. FUS on vagus nerve in I/R+FUS group also reduced neutrophil infiltration compared with the I/R group, I/R+vagotomy group and I/R+vagotomy+FUS group. In contrast, the lack of FUS of vagus nerve in the I/R group induced the opposite effect compared with the sham group. FUS on vagus nerve in I/R-24h+FUS group also reduced infarct size, myocardial fibrosis, apoptosis and oxidative stress, and also preserved left ventricular function compared with the I/R-24h group. In this study, our findings were similar as found in previous studies. Chen *et al*^[8]^ showed that episodes of ventricular arrhythmia in the I/R group were significantly higher than those in the low level vagus nerve stimulation (LL-VNS) group. Episodes of VPCs and VT were significantly reduced in the LL-VNS group during ischemic and reperfusion periods when compared with the I/R group. Zhang *et al*^[9]^ reported that subthreshold vagal stimulation (SVS) without heart rate reduction significantly suppressed I/R induced ventricular arrhythmias and decreased serum concentrations of CRP, IL-6, TNF-*α*, HMGB1 during both the ischemic and the reperfusion periods. Another study^[10]^ showed that the levels of TNF-α and IL-6 increased remarkably in the I/R group, and simultaneously Ach content decreased and a large amount of neutrophil infiltration occurred. The levels of TNF-α and IL-6 in the vagus nerve stimulation group declined and Ach content increased and the neutrophils infiltration decreased clearly. Arimura *et al*^[11]^ found that short-term intravenous electrical vagal nerve stimulation(iVNS) delivered prior to coronary reperfusion markedly reduced infarct size and preserved cardiac function one month after acute myocardial infarction. The bradycardic effect plays an important role in the beneficial effect of iVNS. A clinical study^[12]^ showed that low-level tragus stimulation (LL-TS) reduced myocardial I/R injury in patients with ST segment elevation myocardial infarction(STEMI). This proof-of-concept study raised the possibility that this noninvasive strategy may be used to treat patients with STEMI undergoing primary percutaneous coronary intervention. These above studies proved that electrically stimulation on nerve could attenuate I/R injury. Some previous studies showed that ultrasound could also stimulate the nerve to modulate inflammatory reaction. Chen *et al*^[13]^ showed that the delivery of low-intensity pulsed ultrasound stimulation(LIPUS) on atrial ganglion plexus markedly reduced ventricular episodes, reduced infarct size and improved cardiac function. LIPUS significantly reduced the release of TNF-a, IL-1β and HMGB1. Wasilczuk *et al*^[14]^ found that low-intensity focused ultrasound stimulation of the vagus nerve(uVNS) is a promising new way to attenuate serum TNF-a levels. Multiple applications of uVNS statistically reduced serum TNF-a levels on average by 73% compared with the no stimulation controls.

The process of myocardial I/R injury is complex and includes inflammatory activation. Reperfusion triggers a vigorous inflammatory response, and several signal pathways are activated. Therefore, we suggested that one potential mechanism for the cardiac protective effect against I/R damage might involve initiating the activated kinase/signal transducers and inhibit systemic inflammatory response. Cholinergic transmitter Ach can inhibit systemic inflammatory response syndrome and significantly reduce the release of proinflammatory cytokines. Stimulation of peripheral vagus nerve could activate cholinergic anti-inflammatory pathway.

Our study showed that the α7nAchR protein expression in the I/R, I/R+vagotomy and I/R+vagotomy+FUS groups decreased obviously compared with those in the I/R group, there was no statistical significance between the sham group and the I/R+FUS group. These findings suggest that FUS on vagus nerve could regulate cholinergic anti-inflammatory pathway, as verified by the change of the α7nAchR protein expression. It was equally effective as found in previous studies. Zhang *et al*^[10]^ showed that a large number of deep brown α7nAchR granules were seen in the vagus nerve stimulation group. It proved that the right vagus nerve electric stimulation could activate anti-inflammatory pathway and inhibit the systemic and local inflammatory reaction to relieve myocardial I/R injury. Zhao *et al*^[15]^ indicated that vagal nerve stimulation-mediated anti-inflammatory effect mainly involved the cholinergic anti-inflammatory pathway which was dependent on α7nAChR. Calvillo *et al*^[16]^ showed that vagal stimulation decreased infarct size and inflammatory markers during I/R independent of the heart rate. The anti-inflammatory and antiapoptotic properties of the nicotinic pathway were the primary underlying mechanism.

The present study also found that ultrasound modulated the cholinergic anti-inflammatory pathway to reduce I/R injury required the involvement of vagus nerve. After the vagotomy of the vagal nerves, ultrasound stimulation could not achieve the effect of cardioprotection. Wasilczuk *et al*^[14]^ also found that the anti-inflammatory effect of focused ultrasound was no longer evident when a distal vagotomy was performed. Nuntaphum *et al*^[17]^ showed that vagus nerve stimulation(VNS) required the contralateral efferent vagal activities to fully provide its cardioprotection. It was important to note that most of the current devices implanted in animal and clinical investigations, activate both afferent and efferent pathways^[18-24]^.

In the present study, we found that FUS exhibited cardioprotective effect both in structural and functional terms. In the rodent model of I/R-24h, it showed a significant reduction in infarct size and the infiltration of neutrophils in myocardium in rats treated with FUS. A prior investigation demonstrated that the magnetic vagus nerve stimulation therapy could effectively reduce the infarct size, accompany with reducing inflammatory cytokines secretion and the infiltration of neutrophils in myocardium^[25]^. Preservation of cardiac function was evidenced by greater LVEF and LVFS in rats that received FUS. The improvements in systolic function could be related to the smaller size of the infarct myocardium. These findings are consistent with previous study^[17]^ of vagus nerve stimulation as an attractive method exerts cardioprotection against myocardial I/R injury. We also found that apoptosis were alleviated in rats treated with FUS which may have contributed to the myocardial salvage and improvement of cardiac function. Our data also showed that FUS attenuated oxidative stress as evidenced by the change in the SOD and MDA levels. These findings are consistent with previous study showed that vagus nerve stimulation attenuates myocardial I/R injury by antioxidative stress and antiapoptosis reactions in canines^[26]^.

### Implications for clinical practice

Stimulation on right cervical vagus nerve has been proven to be an effective option for treating acute myocardial I/R injury in animal study. In this study, we suggested that FUS on right vagus nerve might be an effective therapeutic strategy for attenuating myocardium inflammatory reaction and apoptosis, antioxidant, reducing infarct size, preventing ventricular arrhythmias and preserved left ventricular function during acute myocardial I/R period.

## Conclusion

In conclusion, The noninvasive method of ultrasonic neuromodulation using FUS of vagus nerve exerted significant cardioprotection on acute myocardial I/R injury. A potential mechanism of such effects of FUS is associated with cholinergic anti-inflammatory pathway.

## Acknowledgements

None

## Declarations

None

## Funds

This study was supported by grants to Juefei Wu from Guangdong Natural Science Funds for Distinguished Young Scholar (Grant 2016 A030306028), Guangzhou Science and Technology Program (Grant 201506010021), and Foundation of President of Nanfang Hospital (Grant 2020Z006 and 2018Z018), and to Qian Zhang from Shenzhen Fundamental Research Program (Grant JCYJ20220530142602006), and to Weibao Qiu from STI 2030-Major Projects (2021ZD0200401), and National Science Foundation Grants of China (82327805).

## Notes

### Competing Interest Statement

The authors have declared no competing interest.

